# Cryo-EM structure of plant urea transporter DUR3 reveals essential role of C-terminal domain in tetramer assembly and insights into proton-coupled transport

**DOI:** 10.1101/2025.05.20.653754

**Authors:** Yanlin Wang, Marriah N. Green, Huajian Lin, Celina Mazurek, Xiang Lin, Tianming Li, Ruiying Wang, Wenjuan Li, Xiaohui Zhao, Peiqiang Feng, Wolf B. Frommer, Jinru Zhang, Michael M. Wudick, Minrui Fan

**Affiliations:** CAS Center for Excellence in Molecular Plant Sciences, Institute of Plant Physiology and Ecology, Chinese Academy of Sciences, Shanghai 200032, China; Key Laboratory of Plant Carbon Capture, Chinese Academy of Sciences, Shanghai 200032, China; University of Chinese Academy of Sciences, Beijing 101408, China; Heinrich Heine University Düsseldorf, Faculty of Mathematics and Natural Sciences, Institute for Molecular Physiology, 40225 Düsseldorf, Germany; State Key Laboratory of Genetic Engineering, Collaborative Innovation Center of Genetics and Development, Department of Biochemistry and Biophysics, School of Life Sciences, Fudan University, Shanghai 200433, China; Cluster of Excellence on Plant Sciences, Heinrich Heine University Düsseldorf, 40225 Düsseldorf, Germany; Institute for Transformative Biomolecules (WPI-ITbM), Nagoya University, Nagoya 464-8601, Japan

## Abstract

Nitrogen pollution, caused by the overuse of nitrogen-fertilizers such as urea in agricultural production, has become a serious problem that threatens both the environment and human health. Improving the nitrogen use efficiency of crops is an important approach to reduce fertilizer use. DUR3, a high-affinity urea transporter, enables plants to acquire urea from natural or agricultural soils with low urea concentrations and has been proposed as a promising target for engineering crops to improve nitrogen-fertilizer use efficiency. Despite extensive studies for more than 20 years, the structure, substrate recognition and transport mechanism of DUR3 have remained unknown. Here we report the cryo-EM structure of maize DUR3 (ZmDUR3) at 3.41 Å resolution. ZmDUR3 adopts an amino acid-polyamine-organocation (APC) superfamily fold and is structurally distinct from animal facilitative urea transporters. Our structure provides important insights into urea recognition by DUR3 and suggests that a pair of acidic residues is potentially involved in proton coupling during urea transport. Furthermore, our structural and functional studies indicate that DUR3 assembles into a tetramer and tetramer formation is important for its function. Our results pave the way for engineering plant DUR3 protein for applications in sustainable agricultural production.

---

Nitrogen (N), as a primary mineral element, is essential for plant growth and development. Plants can acquire nitrogen from soil in the forms of nitrate, ammonium or urea through their roots (Miller & Cramer, 2005; Kojima *et al*., 2006). The bioavailability of nitrogen in the soil is often limiting plant growth and development. Therefore, every year large amounts of N-fertilizers are being applied to increase crop yield worldwide. Of the N-fertilizers, urea accounts for more than 50% of global use due to its high N content (46%) and competitive price. Although urea is typically converted to ammonium in the soil before absorption, plants can also directly take up urea through high-affinity urea transporters named DUR3, located in the root cell membrane (Kojima *et al*., 2006). Plant DUR3 belongs to the sodium–solute symporter (SSS) family (Liu *et al*., 2003), distinct from the SLC14 family of facilitative urea transporters in animals (Levin & Zhou, 2014). Although SSS family transporters are typically Na^+^-dependent, DUR3 was hypothesized to function as a H^+^/urea symporter (Liu *et al*., 2003). DUR3 has a *K*_m_ in the micromolar range (3–10 μM) and enables plants to acquire urea from natural or agricultural soils with low urea concentrations. The importance of DUR3 transporters in urea acquisition has been demonstrated in multiple plant species, including *Arabidopsis* and crop plants such as rice and maize. Consistent with its role, *DUR3* gene expression in plant roots is upregulated under nitrogen-deficient conditions. Interestingly, *DUR3* is also expressed in other plant tissues, such as senescent leaves, and it has been shown to contribute to nitrogen reallocation and recycling (Liu *et al*., 2003; Kojima *et al*., 2007; Wang *et al*., 2012; Zanin *et al*., 2014; Bohner *et al*., 2015; Liu *et al*., 2015; Beier *et al*., 2019). Due to its physiological importance in plant acquisition and allocation of urea, DUR3 has been proposed as a promising target for engineering crops to improve N-fertilizer use efficiency in agricultural production (Wang *et al*., 2012; Liu *et al*., 2015). Improving nitrogen efficiency is of particular relevance due to the large amounts of energy required for production and the negative impact of nitrogen leaching on groundwater quality and as a result for human health. Despite its importance, the mechanisms by which DUR3 recognizes and transports urea have remained unknown.

To elucidate the structure-function relationship and gain insights into the transport mechanism of DUR3, we set out to determine its structure using cryo-electron microscopy (cryo-EM). After small-scale expression and purification screening of DUR3 proteins from multiple plant species, we selected maize DUR3 (*Zm*DUR3) for its best expression level and peak profile on the gel filtration column (Fig. S1). We determined the structure of *Zm*DUR3 in the presence of 5 mM urea at 3.41 Å resolution (Fig. S2, Table S1). The cryo-EM density indicated that *Zm*DUR3 assembles into a tetramer, with each protomer comprising a transmembrane domain (TMD) and a cytosolic C-terminal domain (CTD) (Fig. 1a–c). Because the subunit A in the *Zm*DUR3 structure exhibited the highest-quality density map, we will focus on it for protomer structural analysis. The TMD consists of 15 transmembrane helices (TM0–14) and adopts an amino acid–polyamine– organocation (APC) superfamily fold with TM1–5 and TM6–10 forming inverted repeats (Fig. 1d). The TMs are connected by loops containing helical elements (α1–4). On the extracellular side, the N-terminal region preceding TM0 interacts with the long loops between TM5 and TM6 (EL3) and TM7 and TM8 (EL4), forming a cap domain on top of the helix bundle composed of TM1, TM2, TM6 and TM7. Notably, two conserved disulfide bonds are involved in stabilizing the N-terminal part and the loop EL3 (Fig. 1d, Fig. S3). The cytosolic CTD consists of two α-helices (α5 and α6) and is connected to the TMD through TM14, which is unique to DUR3 compared to other SSS family transporters, such as the sodium-glucose cotransporter (SGLT), which contains 14 TMs (TM0–13) (Han *et al*., 2022; Niu *et al*., 2022).

**Fig. 1.**
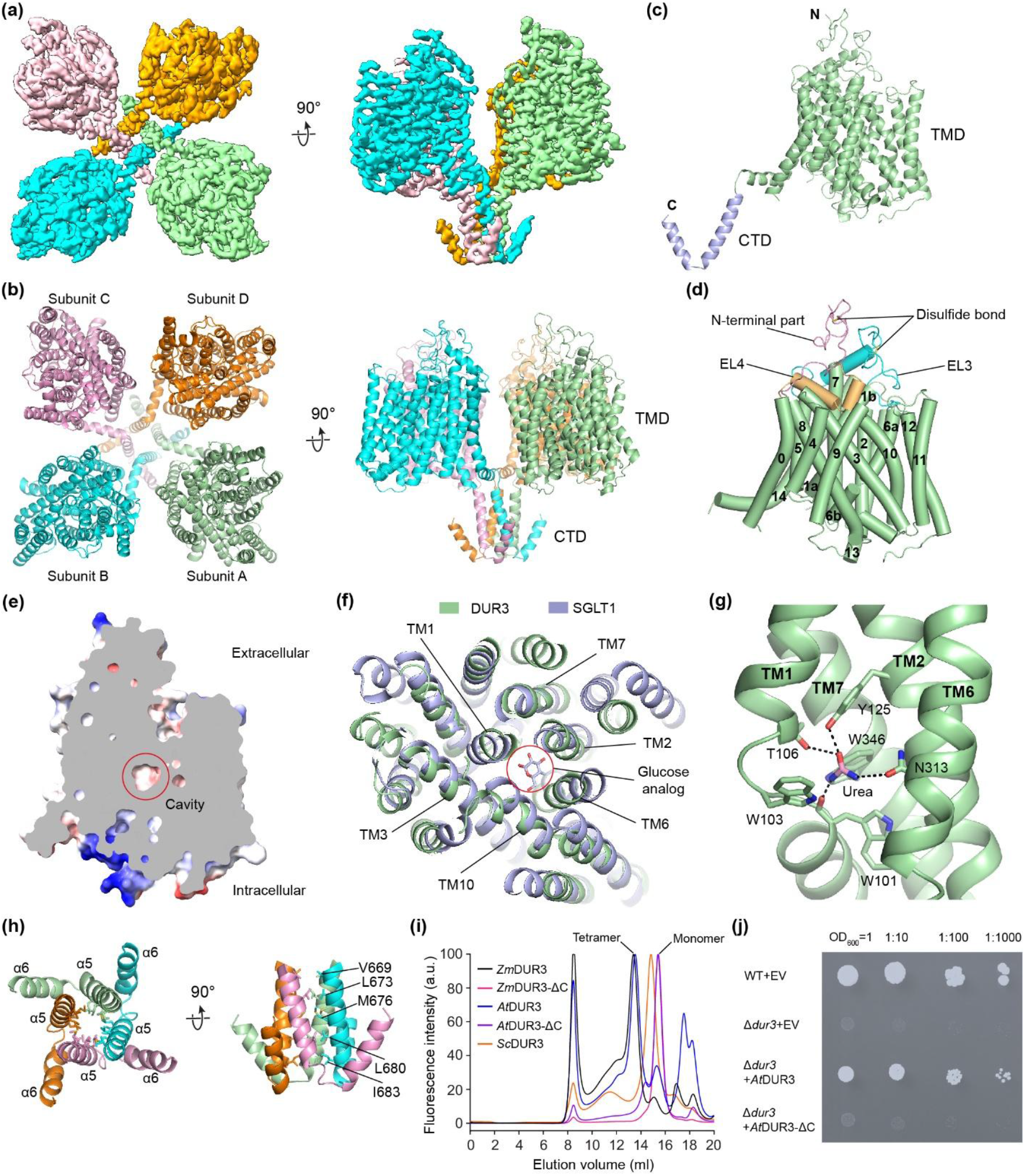
Structure and putative transport mechanism of maize urea transporter *Zm*DUR3. (a) Cryo-EM density map of *Zm*DUR3. (b) Overall structure of *Zm*DUR3. The four protomers are colored palegreen, cyan, pink and orange, respectively. (c) Subunit structure of *Zm*DUR3. The TMD and CTD are colored palegreen and lightblue, respectively. (d) TMD structure in one *Zm*DUR3 subunit. The N-terminal part and the extracellular loops EL3 and EL4 are colored pink, cyan and lightorange, respectively. (e) Sliced view of *Zm*DUR3 in the occluded state. The potential urea-binding cavity is indicated with a red circle. (f) Structural comparison of *Zm*DUR3 and human SGLT1. The glucose analog bound to SGLT1 is shown as sticks. (g) The urea-binding site. The urea-interacting residues are shown as sticks. The hydrogen bonds between urea and surrounding residues are shown as dashed lines. (h) The CTD interactions. (i) Oligomeric analysis of DUR3 using fluorescence-detection size-exclusion chromatography. The full-length and CTD-deleted (ΔC) maize (*Zea mays* L., *Zm*) and *Arabidopsis thaliana* (*At*) DUR3 as well as the full-length yeast (*Saccharomyces cerevisiae, Sc*) DUR3 were characterized. (j) Functional characterization of DUR3 using yeast complementation assay. Wild-type (WT) yeast was used as a positive control, and the Δ*dur3* yeast mutant was used for complementation. EV denotes empty vector.

Our structure captured *Zm*DUR3 in an occluded state, with the substrate translocation pathway closed at both the extracellular and intracellular sides. A cavity is present in the middle of the TMD, which is lined by residues from TM1, TM2, TM3, TM6, TM7 and TM10 (Fig. 1e,f). Because a cavity at a similar position has been identified as the substrate-binding site in human SGLT (Cui *et al*., 2023), which shares the highest structural similarity with *Zm*DUR3 (Z-score=24.5, RMSD=3.6 Å) according to the Dali protein structure comparison server analysis, we suggest that urea may bind to this cavity in *Zm*DUR3. Consistently, we observed a non-protein density in the cavity and tentatively assigned it to urea (Fig. S4a). The urea molecule is sandwiched between two tryptophans (Trp103^TM1^ and Trp346^TM7^), and it also forms hydrogen bond interactions with surrounding residues, including Trp101^TM1^, Thr106^TM1^, Tyr125^TM2^ and Asn313^TM6^ (Fig. 1g). Notably, these six residues are fully conserved in DUR3 across different species (Fig. S3), supporting potential roles in urea binding and transport. The involvement of aromatic residues in urea binding had also been reported in other urea transporters and channels (Levin *et al*., 2012; Strugatsky *et al*., 2013; Levin & Zhou, 2014; Wang *et al*., 2023; Huang *et al*., 2024). Regarding the ion-coupling mechanism of DUR3 transport of urea, our sequence and structural analyses revealed that despite belonging to the SSS family, a key serine involved in Na^+^ coordination in TM8 of canonical SSS family transporters (such as SGLT) is not conserved in DUR3 (Fig. S4b,c). This finding aligns with a previous study suggesting that DUR3 is a proton-coupled urea transporter (Liu *et al*., 2003). Interestingly, we identified a pair of acidic residues – Asp320 from TM6 interacting with Glu410 in TM8 (approximately 3.0-Å distance) – on one side of the substrate-binding cavity (Fig. S4d). This proximity indicates that one of the residues is protonated, similar to observations in chicken acid-sensing ion channel 1 (Jasti *et al*., 2007). Both acidic residues are perfectly conserved in DUR3 across species (Fig. S3). Presumably, changes in the protonation state of Asp320 or Glu410 could affect the interaction between TM6 and TM8 and trigger a conformational change of DUR3. Therefore, we suggest that the Asp320-Glu410 pair may be involved in proton coupling of DUR3 transport of urea.

In the *Zm*DUR3 structure, the tetramer is mainly stabilized by interactions between the CTDs of the four subunits, while the intersubunit interactions in the TMD appear weaker and are potentially mediated by lipid molecules (Fig. 1a,b). The CTDs of *Zm*DUR3 form a four-helix bundle through their α5 helices, with hydrophobic residues pointing to the center and interacting with one another (Fig. 1h). The CTD is relatively conserved in plant DUR3s across different species, but shows considerable variation in yeast DUR3 (Fig. S3). Consistently, fluorescence-detection size-exclusion chromatography revealed that DUR3 proteins from maize and other plants (including *Arabidopsis*, rice and the moss *Physcomitrium patens*) all form tetramers while yeast DUR3 does not. Moreover, deleting the CTD of plant DUR3 led to disassociation of the tetramer (Fig. 1i, Fig. S5). To investigate the potential role of tetramer formation in DUR3 function, we assessed the ability of full-length and CTD-deleted plant DUR3 to rescue the growth of a *dur3*-knock-out yeast mutant (Δ*dur3*) using urea as the sole nitrogen source. Our results demonstrated that full-length *Arabidopsis* DUR3, but not the CTD-deleted version, could rescue the growth of the yeast mutant (Fig. 1j), highlighting the critical role of CTD-mediated tetramer formation in DUR3-mediated transport of urea.

In summary, we have determined the first structure of a plant urea transporter DUR3. Our structural and functional studies suggest that plant DUR3 functions as a tetramer. Additionally, our structure also provides important insights into urea recognition and proton-coupled transport by DUR3. These findings pave the way for engineering DUR3 to enhance N-fertilizer use efficiency in crops.

## Acknowledgements

We thank the State Key Laboratory of Genetic Engineering Cryo-EM Facility of Fudan University for sample imaging. We thank M. Zhang at the Cryo-EM Facility at CAS Center for Excellence in Molecular Plant Sciences for help with sample imaging. We thank N. von Wirén for sharing the yeast strains. We thank A. Plett for her valuable advice on the yeast assay. This work was made possible by support from the CAS Center for Excellence in Molecular Plant Sciences, the Strategic Priority Research Program of Chinese Academy of Sciences (XDB0630101) and the CAS Pioneer Hundred Talents Program to M.F. J.Z. was supported by startup funding from Fudan University. M.M.W. and W.B.F. were supported by the Deutsche Forschungsgemeinschaft (DFG) under Germany’s Excellence Strategy (EXC-2048/1, Project ID 390686111) and the DFG Collaborative Research Center SFB 1208 (Project ID 267205415). Funding was also provided through a Henriette Herz Fellowship awarded to M.N.G.

## Competing interests

None declared.

## Author contributions

MF, MMW and WBF conceived the project. YW and TL carried out biochemical experiments and protein purification. HL processed the cryo-EM data and built and refined the model with MF. MNG and CM performed the functional studies under the supervision of MMW. XL and JZ collected cryo-EM data. RW, WL and XZ contributed to the biochemical experiments. MF, JZ, MMW and WBF analyzed the results. MF wrote the draft manuscript, and MMW, MNG, WBF and PF edited the manuscript.

## Data availability

The cryo-EM density map of *Zm*DUR3 has been deposited in the Electron Microscopy Data Bank and the accession number is EMD-xxxx. The corresponding atomic coordinate has been deposited in the Protein Data Bank and the accession numbers is PDB-xxxx.

## Supporting Information

**Fig. S1** Biochemical characterization of maize DUR3 (*Zm*DUR3).

**Fig. S2** Cryo-EM data processing of *Zm*DUR3.

**Fig. S3** Sequence alignment of DUR3 from different species.

**Fig. S4** Urea-binding site and proton coupling of *Zm*DUR3.

**Fig. S5** CTD is important for tetramer formation of plant DUR3.

**Table S1** Cryo-EM data collection, refinement and validation statistics.

**Methods S1** Materials and methods

**Fig. S1.**
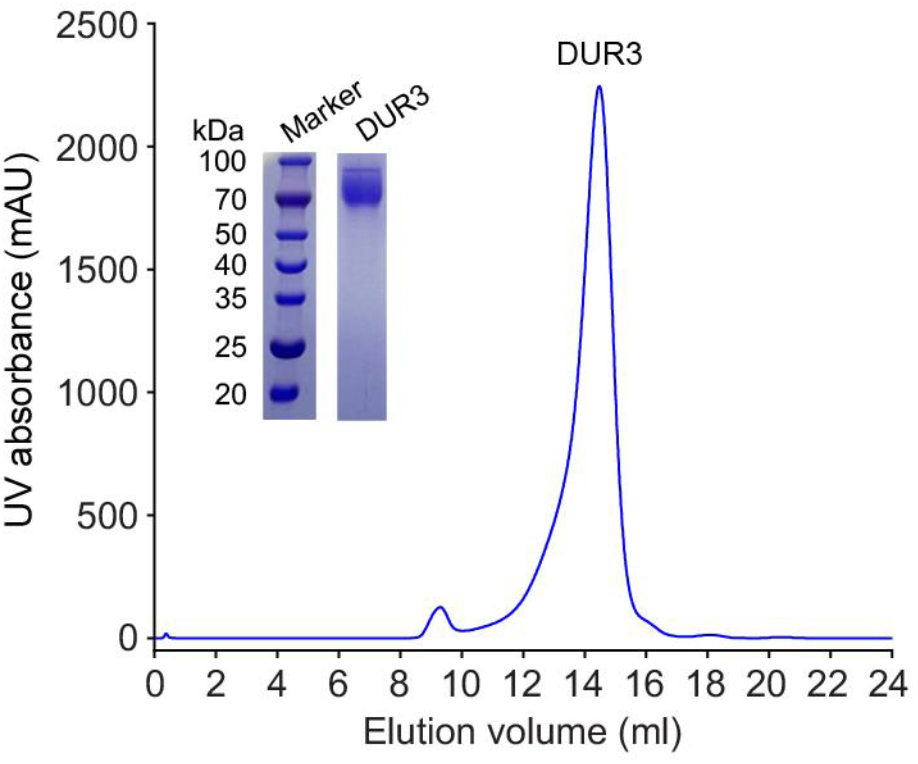
Biochemical characterization of maize DUR3 (*Zm*DUR3). Size-exclusion chromatography profile and SDS-PAGE gel of *Zm*DUR3.

**Fig. S2.**
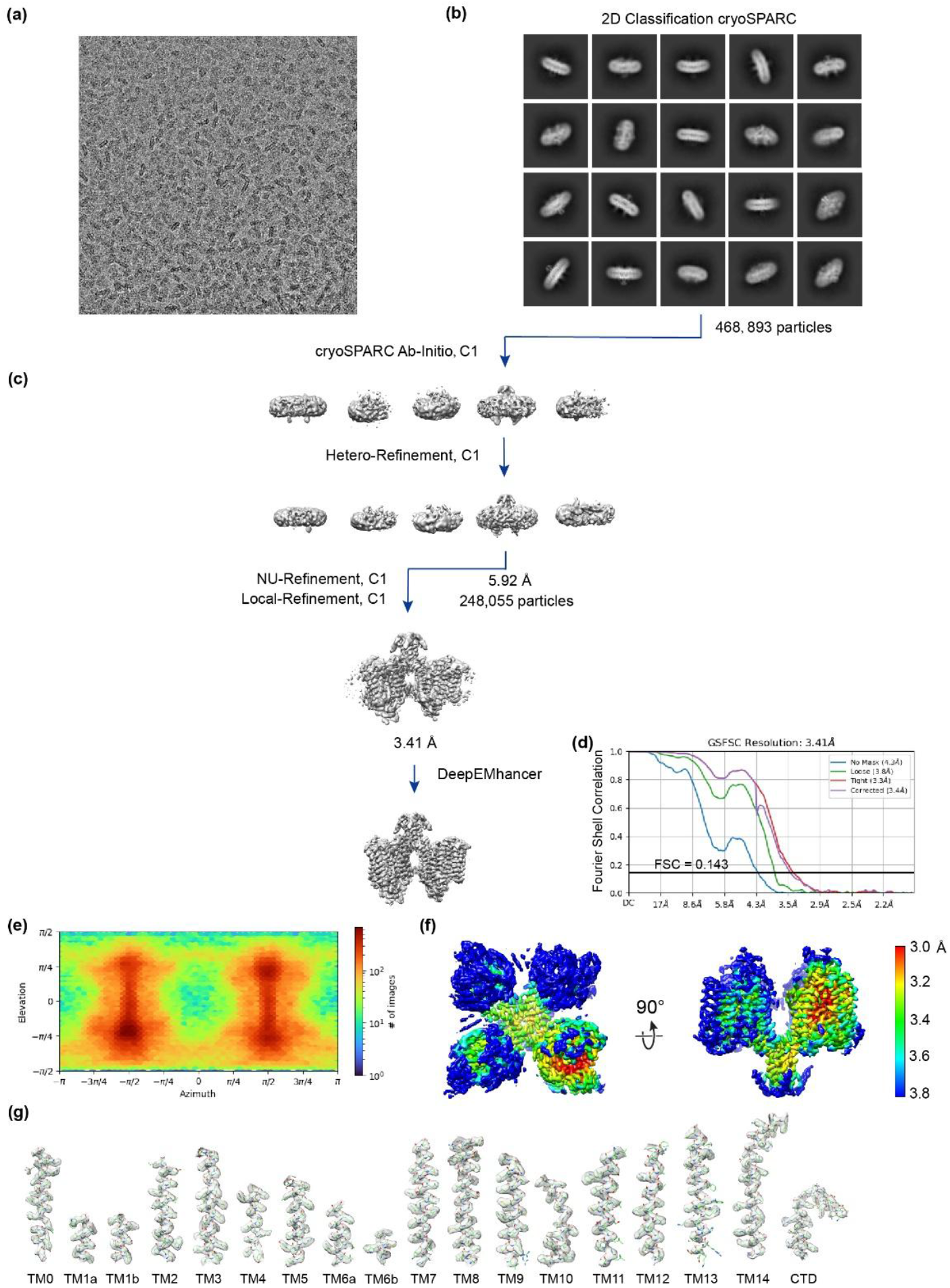
Cryo-EM data processing of *Zm*DUR3. (a) Representative cryo-EM micrograph of *Zm*DUR3. (b) 2D class averages of *Zm*DUR3. (c) Flowchart of data processing of *Zm*DUR3. (d) Gold-standard FSC curves of the final cryo-EM map. (e) Angular distribution of particles for the final reconstruction. (f) Local resolution of the final map. (g) Cryo-EM density maps of *Zm*DUR3.

**Fig. S3.**
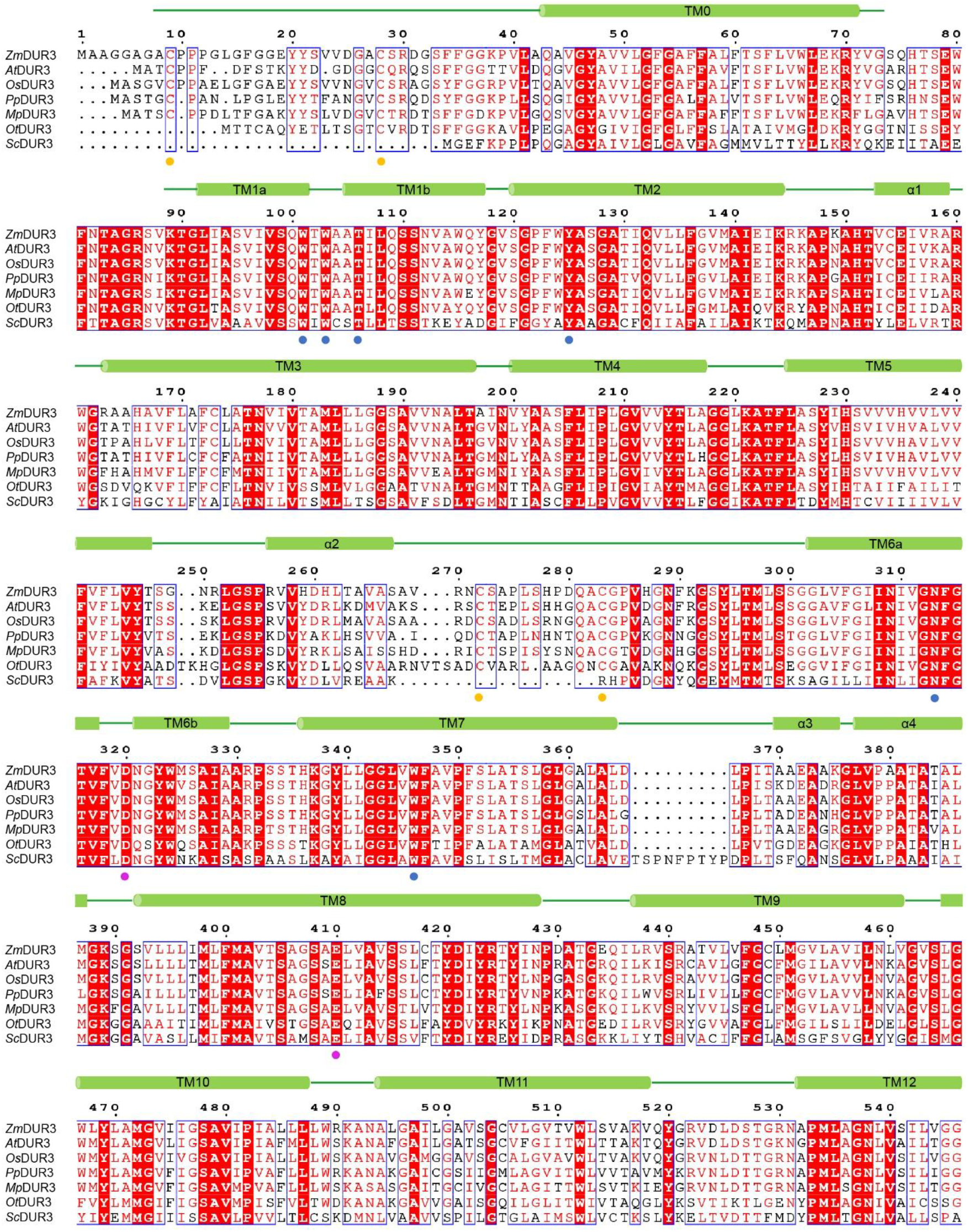
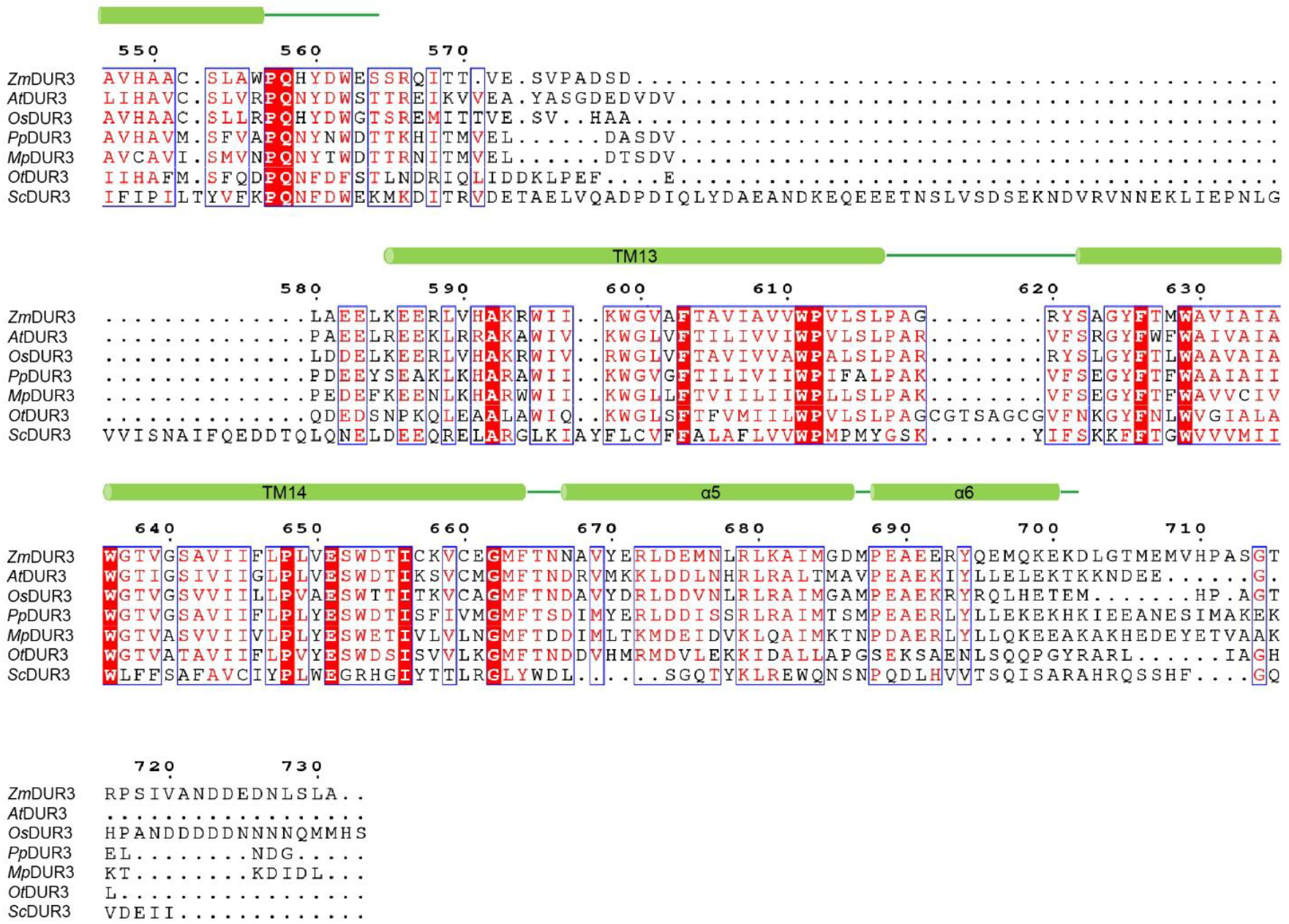
Sequence alignment of DUR3 from different species. The sequences of DUR3 from *Zea mays* L. (*Zm*), *Arabidopsis thaliana* (*At*), *Oryza sativa* (*Os*), *Physcomitrium patens* (*Pp*), *Marchantia polymorpha* (*Mp*), *Ostreococcus tauri* (*Ot*) and *Saccharomyces cerevisiae* (*Sc*) are aligned. The four residues forming disulfide bonds (Cys9-Cys28 and Cys272-Cys283) are indicated with orange dots. The urea-interacting residues are indicated with blue dots. The potential proton-coupling related Asp320-Glu410 pair is indicated with magenta dots.

**Fig. S4.**
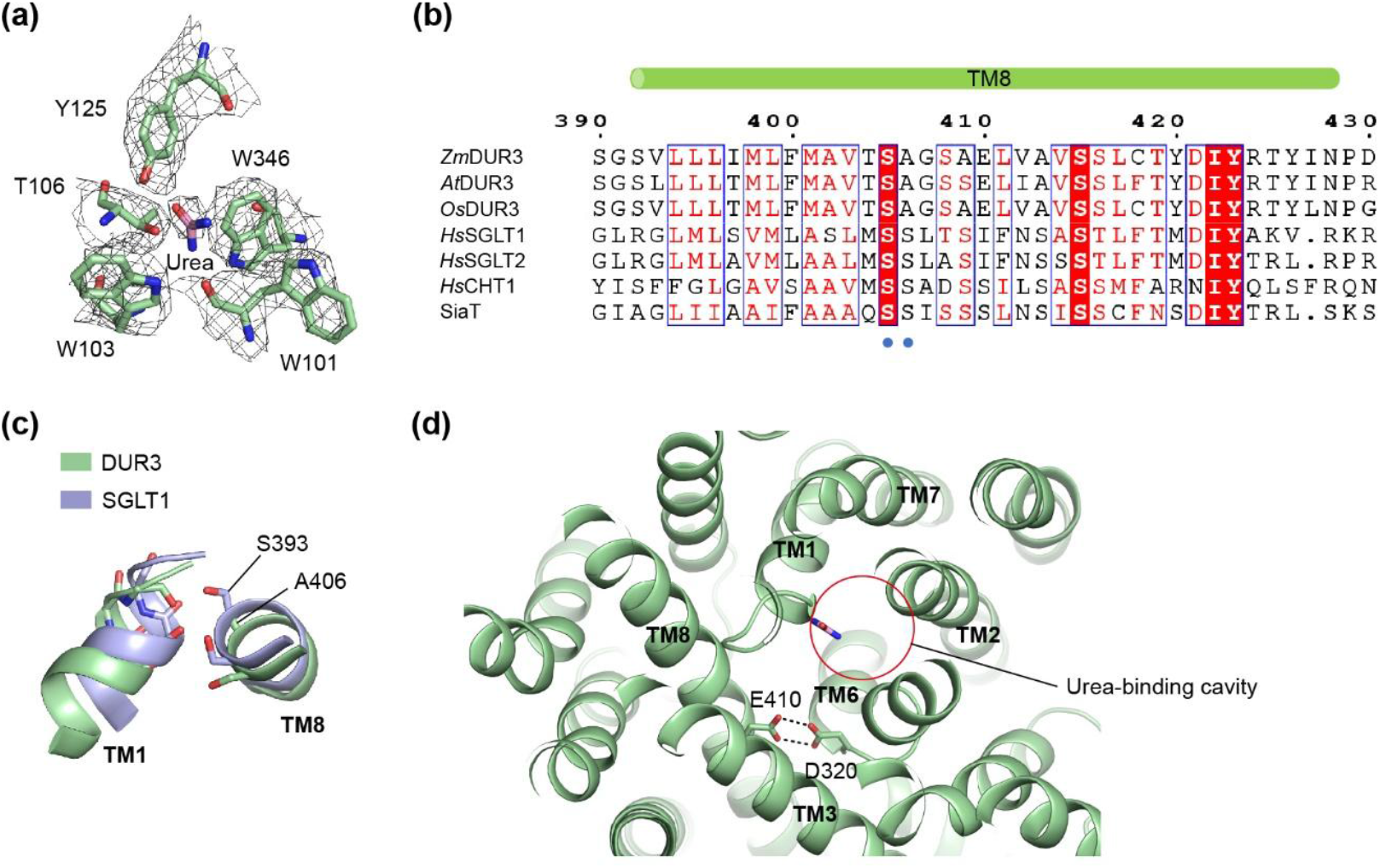
Urea-binding site and proton coupling of *Zm*DUR3. (a) Cryo-EM densities of urea and surrounding residues. The urea and surrounding residues are shown as pink and palegreen sticks, respectively. (b) Sequence alignment of DUR3 and other SSS family transporters. The sequences of DUR3 from maize (*Zm*DUR3), *Arabidopsis* (*At*DUR3) and rice (*Os*DUR3) and the sequences of human glucose transporters SGLT1 and SGLT2 (*Hs*SGLT1 and *Hs*SGLT2), human choline transporter CHT1 (*Hs*CHT1) and the bacterial sialic acid transporter (SiaT from *Proteus mirabilis*) are aligned. The two key serine residues involved in Na^+^ coordination in TM8 of SSS family transporters are indicated with blue dots. (c) A key Na^+^-coordinating serine is not conserved in DUR3 based on structural comparison of *Zm*DUR3 and human SGLT1. Typically, at the canonical Na2 site of SSS family transporters, the side-chain oxygens of two conserved serines in TM8 are involved in coordinating Na^+^, and mutation of either serine would significantly impair transport activity. (d) The Asp320-Glu410 pair at one side of the urea-binding cavity. The distances between the carboxyl oxygens of Asp320 and Glu410 are 2.9 and 3.2 Å, respectively.

**Fig. S5.**
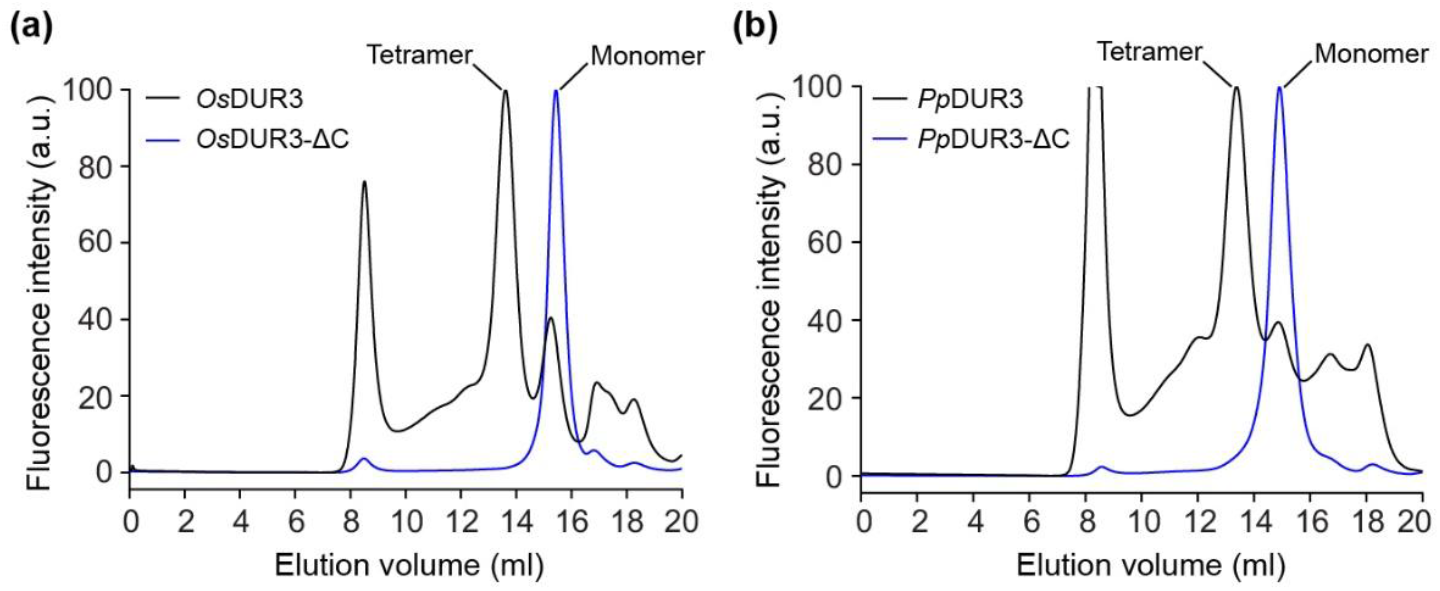
CTD is important for tetramer formation of plant DUR3. (a) Oligomeric analysis of full-length and CTD-deleted (ΔC) rice DUR3 (*Os*DUR3) using fluorescence-detection size-exclusion chromatography (FSEC). (b) Oligomeric analysis of full-length and CTD-deleted (ΔC) *Physcomitrium patens* DUR3 (*Pp*DUR3) using FSEC.

**Table S1.**
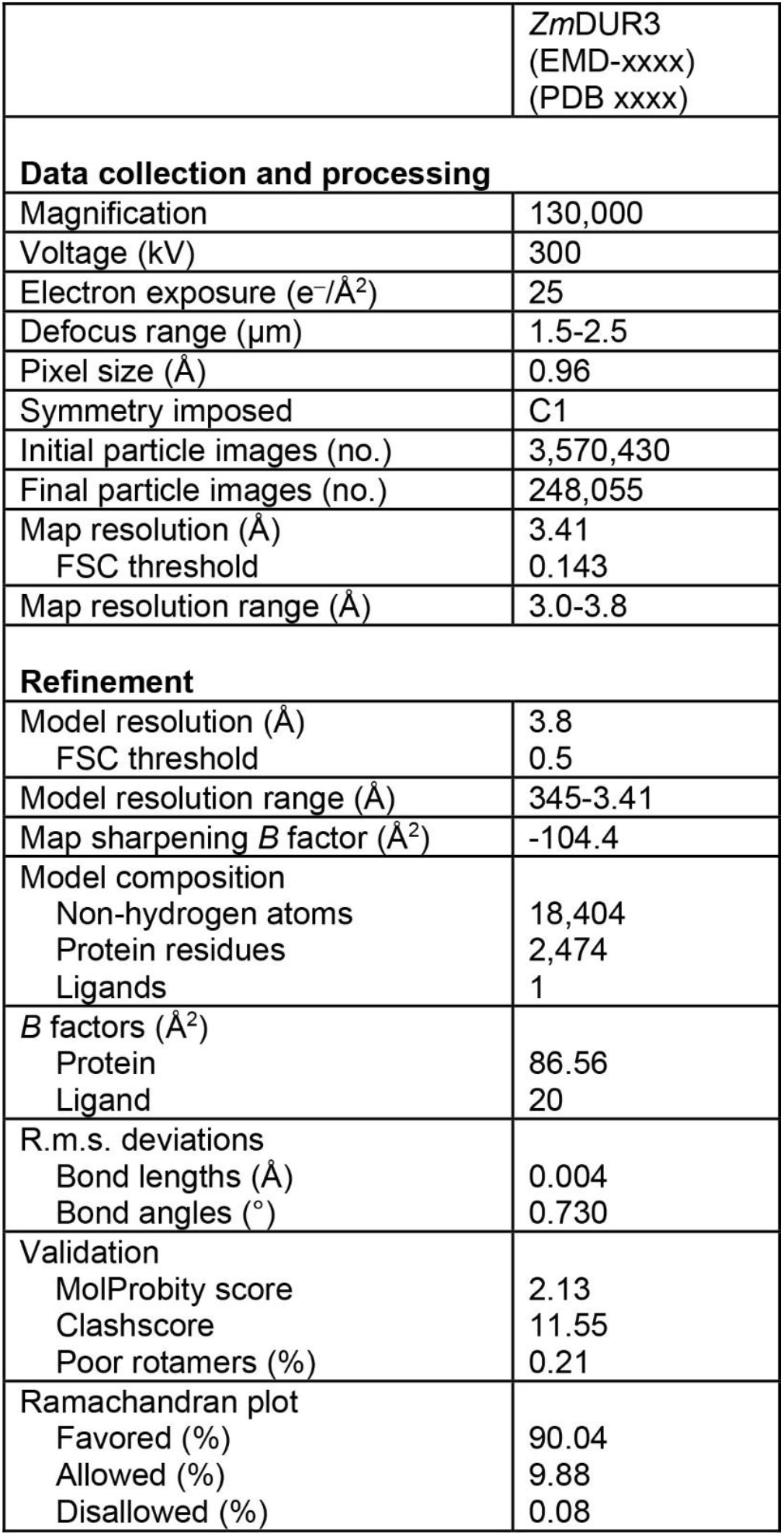
Cryo-EM data collection, refinement and validation statistics.

## Methods S1 Materials and methods

### Protein expression and purification

The coding sequence of DUR3 from *Zea mays* L. with a C-terminal Strep-tag was cloned into a modified BacMam vector. Baculovirus was generated using the Bac-to-Bac system according to established protocols (Goehring *et al*., 2014), and 100 mL of P3 virus was used to infect 1 L of HEK293F cells. After 12 h, 10 mM sodium butyrate was added to boost protein expression, and the cells were cultured at 30 °C for another 48 h before harvesting.

To purify *Zm*DUR3, the cell pellet was resuspended in buffer A (50 mM Tris pH 7.4, 150 mM NaCl and protease inhibitors) and homogenized using a Dounce homogenizer. Crude membranes were obtained by centrifugation at 38,900 *g* for 60 min. The crude membrane fractions were resuspended in buffer A and solubilized with 1.5% lauryl maltose neopentyl glycol (LMNG, Anatrace)/0.2% cholesteryl hemisuccinate (CHS, Anatrace) at 4 °C for 2 h. After centrifugation (38,900 *g*, 50 min), the supernatant was incubated with prewashed Streptactin Beads 4FF (Smart Lifesciences) at 4 °C for 2 h. The slurry was poured into a gravity column (Bio-Rad), and the resin was washed with buffer B (20 mM Tris pH 7.4, 150 mM NaCl and 0.05% glyco-diosgenin (GDN, Anatrace)). The protein was eluted with buffer C (20 mM HEPES pH 7.4, 150 mM NaCl and 0.006% GDN) supplemented with 7.5 mM dethiobiotin. The eluted protein was concentrated and further purified by gel filtration using a Superose 6 Increase column (Cytiva) on a protein purification system (Inscinstech Co., Ltd.). To purify *Zm*DUR3 bound with urea, 5 mM urea was added to the buffers throughout the purification. The sample was concentrated to approximately 15 mg/mL for cryo-EM grid preparation.

### Cryo-EM sample preparation and data collection

Grids (300 mesh, R2/1; Quantifoil, Germany) were glow-discharged using a PELCO easiGlow machine (15 mA, 1 min). Subsequently, 3.5 μL of protein solution was applied to the grids and incubated for 20 sec at 8 °C with 100% humidity. The grids were then blotted with filter paper for 5 sec and plunge-frozen in liquid ethane cooled by liquid nitrogen using a Vitrobot (Vitrobot Mark IV, Thermo Fisher Scientific). The cryo-EM dataset for *Zm*DUR3 was collected using a 300 kV electron microscope (Titan Krios G4, Thermo Fisher Scientific) equipped with a Gatan K3 direct electron detector and EPU software. Data collection was performed at a magnification of approximately 130,000x, yielding a physical pixel size of 0.96 Å. Movies were recorded with a total exposure of 25 electrons/Å^2^, and the defocus was adjusted to range from 1.5 to 2.5 μm. The data collection statistics are summarized in Table S1.

### Cryo-EM data processing

The cryo-EM dataset for *Zm*DUR3 was processed using cryoSPARC (v4.1.2) (Punjani *et al*., 2017). A total of 8,035 movies underwent Patch Motion correction to address beam-induced drift. The contrast transfer function (CTF) parameters of the micrographs were estimated using Patch CTF. Particles were identified using both Blob Picker and Template Picker. Following several rounds of 2D classification, the remaining particles that exhibited clear secondary structure features were selected for ab-initio 3D model generation, applying C1 symmetry during all subsequent 3D reconstructions. The particles were then subjected to Hetero refinement, utilizing the five volumes generated during ab-initio as references. Particles belonging to a class that produced a cryo-EM map with satisfactory resolution were further refined through Non-uniform refinement and Local refinement, resulting in a map with a resolution of 3.41 Å. The image processing workflows are summarized in Fig. S2.

### Model building and refinement

The sharpened cryo-EM map generated by DeepEMhancer (Sanchez-Garcia *et al*., 2021) served as the foundation for model building. An initial structural model predicted by AlphaFold (Jumper *et al*., 2021) was used as the template. Subsequent manual adjustments to local regions were made using COOT (Emsley *et al*., 2010). The models underwent further refinement through real-space refinement in Phenix (Adams *et al*., 2010). The geometry of the refined models was validated using Molprobity (Chen *et al*., 2010). All structural figures were created using PyMOL (Schrödinger, 2024), UCSF Chimera (Pettersen *et al*., 2004) and ChimeraX (Pettersen *et al*., 2021).

### Fluorescence-detection size-exclusion chromatography

The coding sequences of full-length and C-terminal-domain (CTD)-deleted DUR3 from various species were cloned into a modified BacMam vector that produces a target protein fused with C-terminal green fluorescent protein (GFP). The plasmids were transfected into HEK293F cells following the instructions of Hieff Trans® Liposomal Transfection Reagent (40802ES02, Yeason). After culturing at 37 °C for 48 h, the cells were harvested and resuspended in a buffer containing 50 mM Tris (pH 7.4), 150 mM NaCl and protease inhibitors. Subsequently, 1.5% LMNG/CHS was added to extract membrane proteins. After incubation at 4 °C for 2 h, the lysate was spun at 14,000 rpm for 30 min, and the supernatant was loaded onto a Superose 6 Increase column pre-equilibrated with a buffer containing 20 mM Tris (pH 7.4), 150 mM NaCl and 0.0033% LMNG/CHS. The samples were detected by a fluorescence detector (Shimadzu; Ex/Em = 480/510 nm for the GFP signal) with a flow rate of 0.4 mL/min.

### Yeast complementation assay

Complementary DNA (cDNA) of full-length and CTD-truncated (ΔC, deletion of Asn667 to the C-terminus) *Arabidopsis thaliana* DUR3 (*At*DUR3, At5g45380) were cloned into the yeast expression vector pHXT426 (Wieczorke *et al*., 1999) using the Taraka CloneAmp HiFi PCR Premix and the In-Fusion®HD Cloning Kit (Takara Bio). The complementation assay in *Saccharomyces cerevisiae* used Σ 23346c (MAT a, *ura3*) as the wild-type strain and YNVW1 (MATa, *Δura3, Δdur3*) as the *dur3* mutant yeast strain (Liu *et al*., 2003). Yeasts carrying plasmids were cultured in 5 mL CSM-Ura media (MP Biomedicals) containing YNB with ammonium sulfate (MP Biomedicals) and 2 % glucose (ROTH), shaking at 200 rpm for 14–18 h at 30 °C. Under sterile conditions, 1 mL of the yeast culture was pelleted by centrifugation at 3,000 *g* for 5 min, washed 3 times with 1 mL of autoclaved deionized milliQ-H_2_O, centrifuged at 3,000 *g* for 5 min, and the supernatant decanted. Finally, the cells were resuspended in 0.5 mL milliQ-H_2_O. The suspension was diluted with deionized milliQ-H_2_O to OD_600_ 1.0 as the starting point for the dilution series. The complementation assay was carried out on plates containing 2 mM urea (QIAGEN) (added post-autoclaving to prevent degradation) as the sole nitrogen source, supplemented with 2% glucose and YNB without ammonium sulfate or without amino acids (Formedium) and 5 % agar (pH 6). For the complementation assay, dilution series (1, 1:10, 1:100, and 1:1,000) were spotted (3 μL) on the plates, which were wrapped with parafilm to prevent drying and incubated at 30 °C. Growth was monitored and documented at day 7.

## Notes

### Competing Interest Statement

The authors have declared no competing interest.

## References

Beier MP, Fujita T, Sasaki K, Kanno K, Ohashi M, Tamura W, Konishi N, Saito M, Imagawa F, Ishiyama K, et al. 2019. The urea transporter DUR3 contributes to rice production under nitrogen-deficient and field conditions. Physiol. Plant 167(1): 75–89.

Bohner A, Kojima S, Hajirezaei M, Melzer M, von Wiren N. 2015. Urea retranslocation from senescing Arabidopsis leaves is promoted by DUR3-mediated urea retrieval from leaf apoplast. Plant J. 81(3): 377–387.

Cui W, Niu Y, Sun Z, Liu R, Chen L. 2023. Structures of human SGLT in the occluded state reveal conformational changes during sugar transport. Nat. Commun. 14(1): 2920.

Han L, Qu Q, Aydin D, Panova O, Robertson MJ, Xu Y, Dror RO, Skiniotis G, Feng L. 2022. Structure and mechanism of the SGLT family of glucose transporters. Nature 601(7892): 274–279.

Huang SM, Huang ZZ, Liu L, Xiong MY, Zhang C, Cai BY, Wang MW, Cai K, Jia YL, Wang JL, et al. 2024. Structural insights into the mechanisms of urea permeation and distinct inhibition modes of urea transporters. Nat. Commun. 15(1): 10226.

Jasti J, Furukawa H, Gonzales EB, Gouaux E. 2007. Structure of acid-sensing ion channel 1 at 1.9 Å resolution and low pH. Nature 449(7160): 316–323.

Kojima S, Bohner A, Gassert B, Yuan L, von Wiren N. 2007. AtDUR3 represents the major transporter for high-affinity urea transport across the plasma membrane of nitrogen-deficient Arabidopsis roots. Plant J. 52(1): 30–40.

Kojima S, Bohner A, von Wiren N. 2006. Molecular mechanisms of urea transport in plants. J. Membr. Biol. 212(2): 83–91.

Levin EJ, Cao Y, Enkavi G, Quick M, Pan Y, Tajkhorshid E, Zhou M. 2012. Structure and permeation mechanism of a mammalian urea transporter. Proc. Natl. Acad. Sci. USA 109(28): 11194–11199.

Levin EJ, Zhou M. 2014. Structure of urea transporters. In Urea Transporters. Subcellular Biochemistry (eds. Yang, B., Sands, J.), pp. 65–78. Dordrecht: Springer.

Liu GW, Sun AL, Li DQ, Athman A, Gilliham M, Liu LH. 2015. Molecular identification and functional analysis of a maize (Zea mays) DUR3 homolog that transports urea with high affinity. Planta 241(4): 861–874.

Liu LH, Ludewig U, Frommer WB, von Wiren N. 2003. AtDUR3 encodes a new type of high-affinity urea/H+ symporter in Arabidopsis. Plant Cell 15(3): 790–800.

Miller AJ, Cramer MD. 2005. Root nitrogen acquisition and assimilation. Plant Soil 274: 1–36.

Niu Y, Liu R, Guan C, Zhang Y, Chen Z, Hoerer S, Nar H, Chen L. 2022. Structural basis of inhibition of the human SGLT2-MAP17 glucose transporter. Nature 601(7892): 280–284.

Strugatsky D, McNulty R, Munson K, Chen CK, Soltis SM, Sachs G, Luecke H. 2013. Structure of the proton-gated urea channel from the gastric pathogen Helicobacter pylori. Nature 493(7431): 255–258.

Wang C, Zhu WJ, Ding HT, Liu NH, Cao HY, Suo CL, Liu ZK, Zhang Y, Sun ML, Fu HH, et al. 2023. Structural and molecular basis for urea recognition by Prochlorococcus. J. Biol. Chem. 299(8): 104958.

Wang WH, Kohler B, Cao FQ, Liu GW, Gong YY, Sheng S, Song QC, Cheng XY, Garnett T, Okamoto M, et al. 2012. Rice DUR3 mediates high-affinity urea transport and plays an effective role in improvement of urea acquisition and utilization when expressed in Arabidopsis. New Phytol. 193(2): 432–444.

Zanin L, Tomasi N, Wirdnam C, Meier S, Komarova NY, Mimmo T, Cesco S, Rentsch D, Pinton R. 2014. Isolation and functional characterization of a high affinity urea transporter from roots of Zea mays. BMC Plant Biol. 14: 222

## Supplementary references

Adams PD, Afonine PV, Bunkoczi G, Chen VB, Davis IW, Echols N, Headd JJ, Hung LW, Kapral GJ, Grosse-Kunstleve RW, et al. 2010. PHENIX: a comprehensive Python-based system for macromolecular structure solution. Acta. Crystallogr. D Biol. Crystallogr. 66(Pt 2): 213–221.

Chen VB, Arendall WB, 3rd, Headd JJ, Keedy DA, Immormino RM, Kapral GJ, Murray LW, Richardson JS, Richardson DC. 2010. MolProbity: all-atom structure validation for macromolecular crystallography. Acta. Crystallogr. D Biol. Crystallogr. 66(Pt 1): 12–21.

Emsley P, Lohkamp B, Scott WG, Cowtan K. 2010. Features and development of Coot. Acta. Crystallogr. D Biol. Crystallogr. 66(Pt 4): 486–501.

Goehring A, Lee CH, Wang KH, Michel JC, Claxton DP, Baconguis I, Althoff T, Fischer S, Garcia KC, Gouaux E. 2014. Screening and large-scale expression of membrane proteins in mammalian cells for structural studies. Nat. Protoc. 9(11): 2574–2585.

Jumper J, Evans R, Pritzel A, Green T, Figurnov M, Ronneberger O, Tunyasuvunakool K, Bates R, Zidek A, Potapenko A, et al. 2021. Highly accurate protein structure prediction with AlphaFold. Nature 596(7873): 583–589.

Pettersen EF, Goddard TD, Huang CC, Couch GS, Greenblatt DM, Meng EC, Ferrin TE. 2004. UCSF Chimera – a visualization system for exploratory research and analysis. J. Comput. Chem. 25(13): 1605–1612.

Pettersen EF, Goddard TD, Huang CC, Meng EC, Couch GS, Croll TI, Morris JH, Ferrin TE. 2021. UCSF ChimeraX: Structure visualization for researchers, educators, and developers. Protein Sci. 30(1): 70–82.

Punjani A, Rubinstein JL, Fleet DJ, Brubaker MA. 2017. cryoSPARC: algorithms for rapid unsupervised cryo-EM structure determination. Nat. Methods 14(3): 290–296.

Sanchez-Garcia R, Gomez-Blanco J, Cuervo A, Carazo JM, Sorzano COS, Vargas J. 2021. DeepEMhancer: a deep learning solution for cryo-EM volume post-processing. Commun. Biol. 4(1): 874.

Wieczorke R, Krampe S, Weierstall T, Freidel K, Hollenberg CP, Boles E. 1999. Concurrent knock-out of at least 20 transporter genes is required to block uptake of hexoses in Saccharomyces cerevisiae. FEBS Lett. 464(3): 123–128.

